# Fibronectin binding and non-canonical transcription factor signature distinguishes lung and bone marrow megakaryocytes

**DOI:** 10.64898/2026.11.23.767589

**Authors:** Gabriel A. Leung, Maksym Jopek, Mark Jones, Haley Benzon, Isaac Ostler, Jesse Rowley, Matthew T. Rondina, Aaron Petrey, Elizabeth A. Middleton

## Abstract

Recent evidence has demonstrated the unique morphology and function of lung megakaryocytes (MKs), yet the source of the differences between lung and bone marrow (BM) MKs remains unclear. While BM MKs are large, polyploid, BM-resident cells primarily known as platelet producing cells, lung MKs are smaller, less polyploid, and known for immunomodulation. To better understand the factors underlying the differences in lung and BM MKs, we developed a flow cytometry method for precise characterization and isolation of MKs using leukocyte exclusion and examination of hematopoietic and canonical MK markers. When MK-leukocyte aggregates are excluded, a single femur+tibia contains approximately 1,493 MKs and a single lung contains approximately 4,541 MKs. Enrichment via positive selection efficiently removes unwanted cells, increasing the percentage of MKs amongst live lung cells by 10.29-fold and the percentage of MKs amongst live BM cells by 29.09-fold. Sorted MKs were viable and spread onto fibrinogen. Gene expression analysis demonstrated that lung MKs increase immune pathway gene expression, including TNFRSF1B. Transcription factor (TF) and cell cycle gene expression decreased in lung MKs compared to BM MKs, including reduced expression of GATA1 and CDC6, possibly explaining observations of low ploidy and size in lung MKs, while expression of the TF ETS2 increased. Lung MKs also express increased ITGA4 compared to BM MKs, abundant surface integrin α4 protein, and increased surface integrin β7 protein compared to BM MKs. Additionally, lung MKs bind significantly more fibronectin *ex vivo* then BM MKs. These data demonstrate for the first time that lung MKs have a unique TF signature and suggests integrin α4β7 as a relevant receptor for lung MK function.

**Key Points:** Fibronectin binding and altered GATA1, ETS2, and CDC6 expression characterizes lung megakaryocytes

## Introduction

Megakaryocytes (MKs) have been reported in the lungs since nearly their first identification in the bone marrow (BM) and recent reports have led to their re-examination via flow cytometry and transcriptomics^1–4^. Multiple groups have observed their distinct morphology: their small size, low ploidy, and immunomodulatory function; yet the origin of differences between lung and BM MKs has yet to be determined^5–7^. Lung MKs have been demonstrated to present antigen to T-cells, and produce platelets under baseline and inflammatory conditions^5, 6, 8^. Simultaneously, BM MKs have been recently reported to perform novel functions, participating in niche regulation and immune function, and notably, distinct functions correlated with distinct morphology^9–15^. These data suggest that distinct morphology may denote distinct functions for lung MKs and that MK influences on health and disease may yet to be uncovered.

Studies of MKs are challenged by heterogeneity due to the presence of leukocytes, which can aggregate with MKs or form platelet-leukocyte aggregates (PLAs)^5, 6, 15–17^. Hematopoietic progenitors may also contribute to heterogeneity, as megakaryocyte progenitors (MkPs) and MKs are both present in the BM^18, 19^. Flow cytometric analysis of MKs is typically based on size, DNA content, integrin α2b expression (CD41a), and GP1b-V-IX (CD42) expression^10, 20–24^. This data is often presented without parent gates, such as singlet, viability, CD45, lineage/leukocyte, or progenitor gates, making replication of published MK data challenging. Thus, flow cytometric MK analysis would benefit from further methodological refinement.

Despite many reports describing the unique morphology and function of lung MKs, to our knowledge, no data has been presented to explain the source of these distinct characteristics. In order to examine underlying transcriptional differences that may drive unique lung MK characteristics, we have developed a high-resolution method for MK characterization, isolation, and transcriptional analysis. We hypothesize that lung-resident MKs express unique cell cycle and transcription factor (TFs) genes, leading to distinct morphology and function. Our data demonstrate that lung MKs express canonical MK surface markers, while being smaller, lower ploidy, and expressing a unique TF signature compared to BM MKs. Finally, we have demonstrated unique lung MK function, showing that they have a greater ability to bind fibronectin compared to BM MKs.

## Methods

### Animals

All mice were bred and housed in the University of Utah AAALAC certified viviaria on a rotating 12-hour light/dark cycle, with access to food and water. 8-16 week old animals were used for experiments, regardless of sex. All procedures were approved by the University of Utah Institutional Animal Care and Use Committee (IACUC). Animals were euthanized by C0_2_ asphyxiation, followed by cervical dislocation.

### Tissue processing

After euthanasia, mouse femur, tibia, and lungs were dissected. BM was centrifugated from femur and tibia. Lung tissue was minced with scissors then digested with 3.6 mg/mL collagenase (Worthington #CLS4) for 30 minutes at 37°C, and homogenized into single cell suspension using 16-gauge and 18-gauge needles.

### MK Flow Cytometry Surface Staining

BM or lung cells were stained with a staining cocktail containing Fc block (anti-CD16/32), leukocyte-specific antibodies (Supplementary Table 1/2), MK-specific antibodies (Supplementary Table 3 and/or 5), and 50 ng/mL DAPI in 5 mM EDTA+1x PBS. Primary staining using biotinylated antibodies and secondary incubation with labeled streptavidin were both performed at 4°C for 20 minutes.

### FACS Isolation

After surface staining and magnetic enrichment, cells were resuspended to 5-10x10^6^ cells/mL, then cells were sorted using 5-laser Cytek Aurora CS system (Cytek Biosciences) at 1,000-2,000 events/second. 10,000 cells were sorted into 0.45 mL of culture media (DMEM F12, 10% FBS, 100 U/mL Pen-Strep, 50 ng/mL TPO).

### Ploidy Analysis

Cells were stained for viability using Zombie Violet (Biolegend, catalog #423113), then surface markers. Cells were fixed with 1x FACS Lyse (BD, catalog #349202) for 10 minutes at room temperature, then DNA was stained with 0.1 mg/mL propidium iodide in 0.05% Triton X-100 for 20 minutes on ice. Cells were washed and resuspended in 5 mM EDTA+1x PBS for analysis.

### Fibronectin Binding

Lung and BM cells were stained for surface markers, then incubated with 10 µg/mL labeled fibronectin for 1 hour at 37°C, then cells were washed with 5 mM EDTA+1x PBS, pelleted at 400x G for 5 minutes, decanted, and resuspended in 5 mM EDTA+1x PBS for analysis.

### Quantification and Statistical Analysis

Experiment number and *n* are described in each figure legend. Statistical significance was determined by unpaired t-test using Graphpad Prism.

### RNA-seq Library Prep, and bulk RNA-sequencing

Libraries were prepped using the NEBNext Ultra II Directional RNA Library Prep Kit for Illumina (NEB #E7760S) and NEBNext Poly(A) mRNA Magnetic Isolation Module (NEB #E7490S). Libraries were sequencing using the Element Biosciences Aviti sequencer.

## Results

### Spectral and imaging flow cytometry identify lung MKs using canonical markers and morphology

Due to the potential for MK-leukocyte aggregation in lung MK analysis we examined leukocyte marker expression with CD41a+ cells. We hypothesized that leukocyte marker expression coincident with CD41a+ cells was likely indicative of either PLAs or MK-leukocyte aggregation. Published reports of PLAs commonly demonstrate aggregation with monocytes, neutrophils, and T-cells, as well as the pan-hematopoietic marker, CD45^25–27^. Similarly, MK interactions with B-cells, macrophages, and osteoblasts have been reported, but are not included in analysis of platelet/MK aggregation^28–30^. In addition to commonly profiled leukocytes in PLA analysis, we also examined expression of B-cell, macrophage, mast cell, and erythrocyte markers coincident with platelet markers (Supplementary Table 1) to gain a broader view of leukocyte aggregation with MKs. Our flow cytometric analysis of murine lung detected substantial lymphocyte aggregation, the B-cell marker B220 being detected with 52.2% of CD41a+ cells, and T-cell CD3 was detected with 2.9% of CD41a+ cells.

Additionally, the myeloid marker CD11b was detected with 19.5% of lung CD41a+ cells, and the presence of macrophages (F4/80), monocytes (Ly6c), and neutrophils (Ly6g) were also detected. (Sfig1A,C). We also examined developmental heterogeneity by measuring expression of CD117 (c-kit), which broadly labels hematopoietic progenitors, including MkPs. Of all live, CD45+ CD41a+ lung cells, 5.33% also express CD117 (Sfig1B,C), suggesting that lung-resident hematopoietic progenitors also exist.

Based on this data we developed a panel of platelet and MK surface markers (Table 3), leukocyte markers (Table 2), and a gating strategy to specifically identify MKs and hematopoietic progenitors. Our gating strategy pre-gated on size, singlets, live cells, and CD45+ cells (Fig1A). Nearly 60% of live lung cells express high levels of CD45, including CD41a+ cells (Sfig2A). To minimize leukocyte contamination, we selected eight leukocyte markers to be excluded (Table 2, Fig1B). A fraction of lung CD41a+ cells express the progenitor marker CD117, therefore we separated presumed progenitors (lung CD117+ MKs: CD45^+^lineage^-^ CD117^+^CD41a^+^) from CD117-MKs (Fig1C). Lung CD117-MKs (lung MKs: CD45^+^lineage^-^CD117^-^ CD41a^+^CD42c^+^) express the canonical MK markers, CD41a and CD42c (Fig1D). Mature MKs are polyploid, and can reach DNA contents as high as 128N^31^. Comparatively, CD45+ leukocytes are 2N, therefore we compared the DNA staining of lung MKs to CD45+ lung leukocytes. DNA staining of lung MKs revealed a range of DNA content, indicating the presence of polyploid nuclei (Fig1E).

MKs are rare cells with reports quantifying BM MKs at 0.05% of total BM^32^. To count MKs we used bead-based quantification, finding that a single murine lung contained 4,541 (0.022% of live) MKs and 2,698 (0.016% of live) CD117+ MKs (Fig1F-G). To confirm lung MK identity, we used imaging flow cytometry and examined MK morphology within our gating strategy. We gated on CD45^+^CD41a^+^CD42c^+^lineage^-^CD117^-^ cells, then examined morphology, CD41a/CD42c cell surface expression, and DNA staining (Sfig3A-C). Using this strategy, we were able to identify nucleated lung MKs expressing CD41a and CD42c. Surface expression of CD41a and CD42c demonstrates that these MKs are not PLAs (Fig1H).

We demonstrate here by flow cytometry that polyploid CD117-CD41a+ CD42c+ MKs are present within the lungs. Similarly, imaging flow cytometry reveals lung MKs are not PLAs, as CD41a and CD42c expression is distributed across the cell membrane, while simultaneously revealing the morphological similarity of lung MKs with BM MKs.

### CD41a+ BM cells have developmental heterogeneity

Like the lung, the BM may also aggregate with leukocytes, therefore, we profiled leukocyte marker detection with BM CD41a+ cells, as in the lung. CD11b was detected with 31.1% of BM CD41a+ cells, as well as F4/80, Ly6c, and Ly6g (Sfig1D,F). B220 was detected with 39.2% and CD3 with 0.3% of BM CD41a+ cells. To determine the level of developmental heterogeneity within BM CD41a+ cells we examined CD117 expression using flow cytometry, revealing that 25.02% of live, CD45+ CD41a+ BM cells also express CD117 (Sfig1E,F).

Having determined that leukocytes aggregate with MKs in the BM as in the lung, we employed the same gating strategy and pre-gated on size, singlets, live cells, CD45+ cells, then excluded lineage (Fig2A,B). The lineage-BM can be divided into either CD117+ progenitors or CD117-cells, which differ in size, CD45 expression, and autofluorescence (Fig2C, SFig2B). BM MkPs (CD45^+^ lineage^-^CD117^+^Sca1^-^CD150^+^CD41a^+^) have been previously defined, residing within the CD117^+^Sca1^-^fraction, and we have integrated their definition into our gating strategy^19^. BM CD117-MKs (BM MKs: CD45^+^lineage^-^CD117^-^CD41a^+^CD42c^+^) are highly polyploid. DNA staining revealed a ploidy range between 2N-32N+, with the majority of BM MKs having 4N-8N nuclei (Fig2D-E). MK quantification revealed that a single femur+tibia contains on average 1,493 (0.0055% of live) BM MKs and 2,707 MkPs (0.0113% of live) (Fig2F-G).

Imaging flow cytometry analysis of BM MKs confirmed established morphological features of BM MKs, revealing large, polyploid cells expressing cell surface CD41a and CD42c expression. Notably, this population was morphologically diverse in size, supporting reports of BM MKs with diverse morphology and function^9^ (Fig2H). MkP images were also collected, demonstrating prominent CD117 expression with CD41a and CD42c expression (Sfig3D).

Our methodology for MK characterization supports existing data on BM MK morphology and ploidy, and also finds that after exclusion of MkPs and MKs complexed with leukocytes, counts of murine BM MKs are less than reported MK counts^32^.

### CD41a based magnetic enrichment and isolation of lung and BM MKs

Having established a gating strategy for lung and BM MK characterization, we next sought to efficiently isolate these populations using a positive-selection strategy for pre-enrichment based on CD41a labeling, similar to a previous report (Fig3A)^33^. To determine the effectiveness of this strategy in enriching for MKs, we compared the frequency of MKs in whole lung or BM to enriched lung or BM. After enrichment, BM MKs comprised 0.16% of live BM cells compared to 0.0055% of live BM cells, representing a 29.09-fold enrichment; similarly, lung MKs comprised 0.239% of live cells compared to 0.022% representing a 10.29-fold enrichment (Fig3B,C). FACS isolation of fragile cells such as MKs can induce apoptosis, however post-sort viability analysis revealed that lung MKs were highly viable, averaging 95.16% live, while BM MKs had slightly lower viability, averaging 84.9% live (Fig3D,E). We next decided to examine their ability to spread on fibrinogen, one of the ligands of CD41a. Confocal imaging showed large MKs with a single multilobulated nucleus spread on fibrinogen with prominent staining for GP1b and actin (Fig3F). BM MkPs were similarly able to spread onto fibrinogen coated slides (SFig4).

### Lung MKs transcribe core MK genes while also expressing transcriptional patterns associated with MK immaturity

Our examination of DNA content had revealed that lung MKs were markedly lower ploidy as compared to BM, suggestive of decreased maturity, leading us to hypothesize that the lung MKs express an immature transcriptional MK program. Using our positive-selection and FACS isolation strategy we sorted 30,499 lung MKs and 14,232 BM MKs, then performed RNA-seq expression analysis. Despite both sources of MKs expressing canonical markers, principal component analysis showed that lung and BM MK transcriptomes clustered according to tissue of origin. (Fig4A). Pathway analysis of biological processes enriched for increased gene expression supported previously published data describing lung MKs as immunomodulatory cells, with top upregulated pathways including “immune effector process” and “regulation of leukocyte activation” (Fig4B). Cell cycle pathways including “chromosome organization”, “organelle fission”, and “nuclear division” were amongst the top biological processes enriched for decreased gene expression, similar to recent reports (Fig4C)^5, 7^. To examine reports of reduced maturity, we examined core MK genes such as ITGA2B, ITGB3, GP1BA, GP6, ITGA4, and PF4 and found that expression of ITGA2B, ITGB3, and PF4 were decreased in lung compared to BM MKs^5, 6^. Expression of GP1BA and GP1BB are also reduced, although they are of lower abundance than ITGA2B, typically found to increase with greater ploidy (Fig4E)^23, 24^. Conversely, ITGA4 expression, typically associated with MkPs and immature MKs, was found to be increased in lung MKs (Figure 4D)^34, 35^.

**Figure 1:**
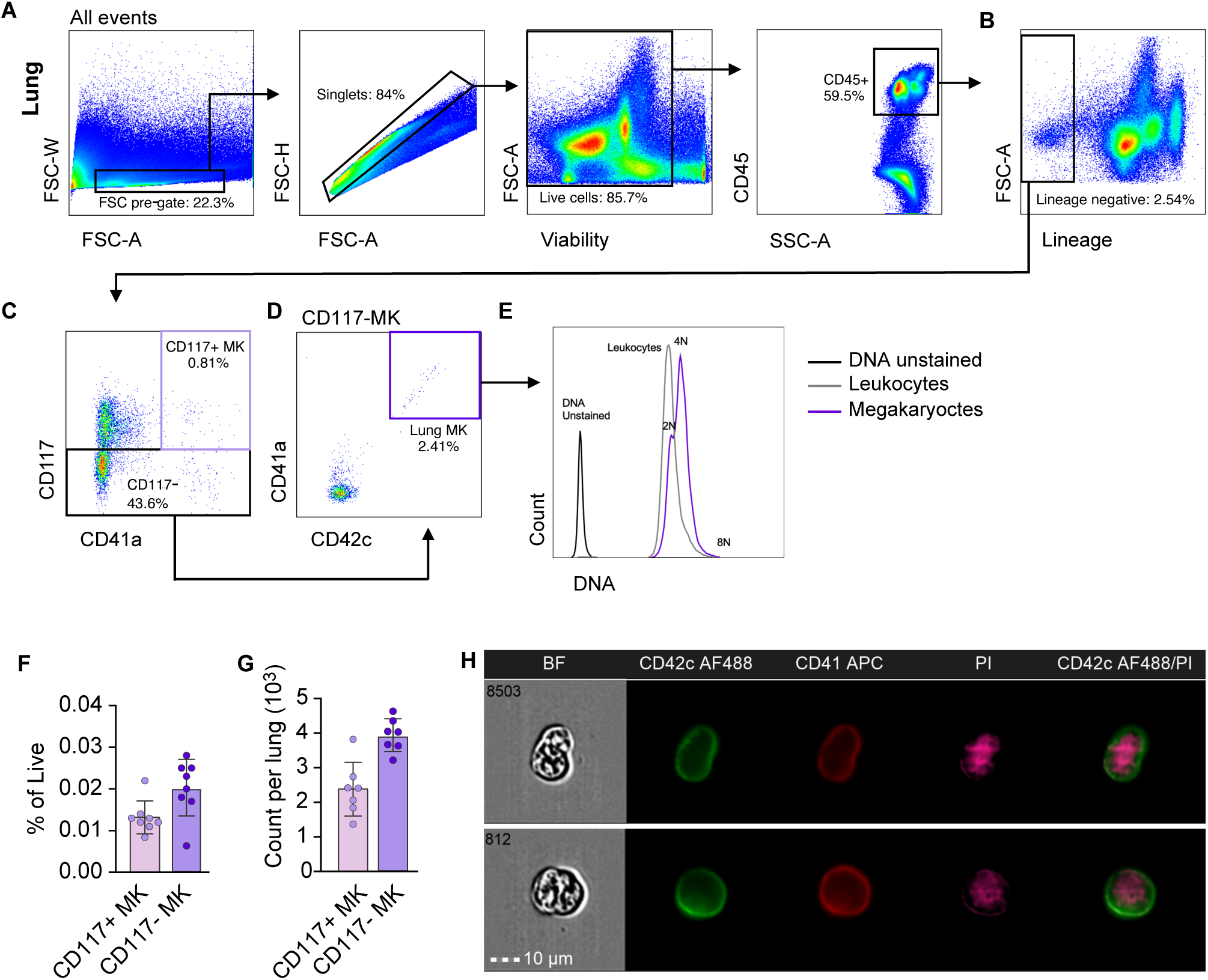
Spectral and imaging flow cytometry identify lung MKs using canonical markers and morphology: (A-D) Representative FACS plots of lung MK gating. (E) Representative DNA staining of lung MKs, lung leukocytes, and DNA staining negative control. (F) Lung MK % of live, n=16 animals, 3 representative experiments. (G) Bead based lung MK quantification, n=14 animals, 3 representative experiments. (H) Representative images from imaging flow cytometry analysis of nucleated live, CD45+ lineage-CD117-CD41a+ CD42c+ cells.

**Figure 2.**
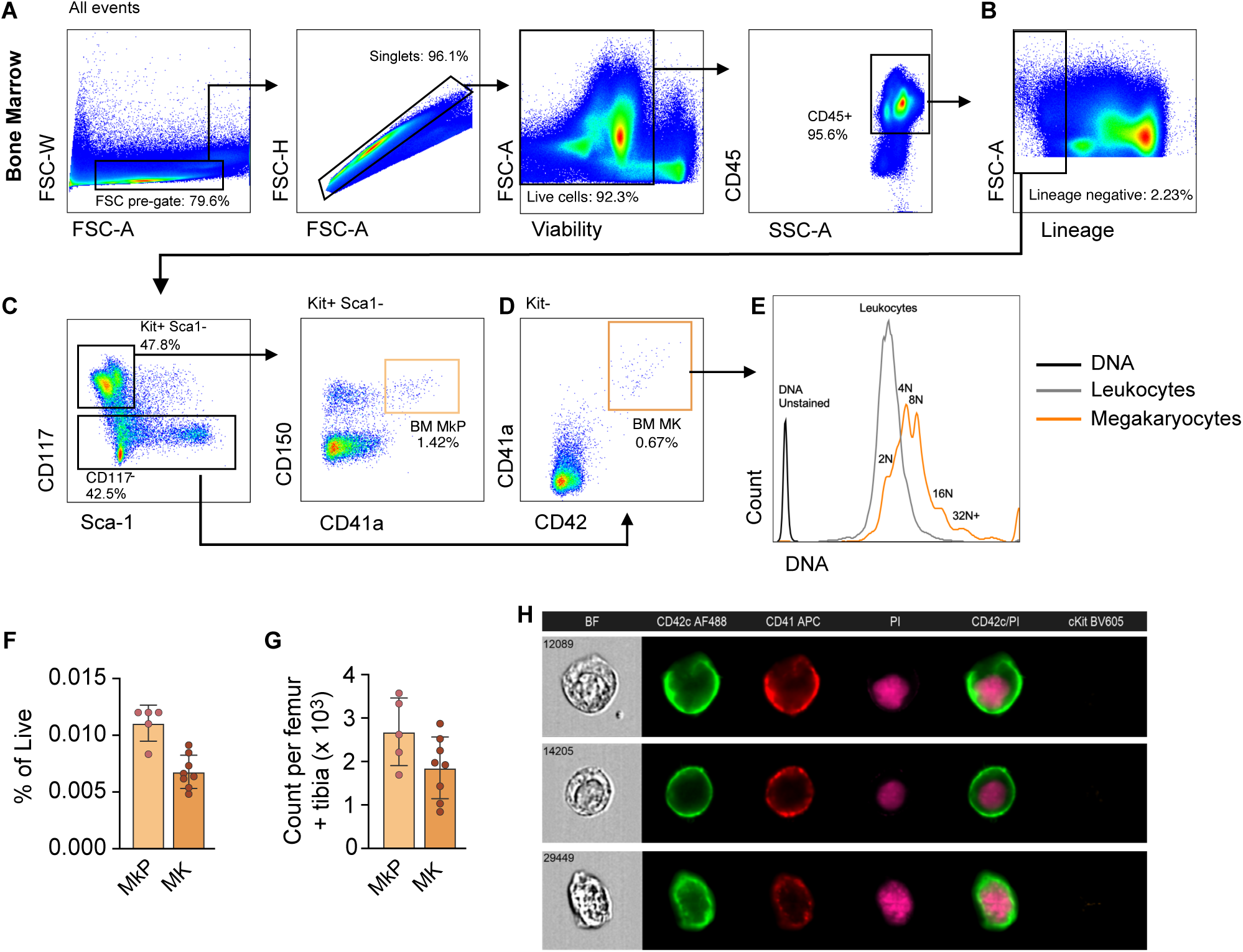
CD41+ BM cells have developmental heterogeneity: (A-D) Representative FACS plots of BM MK gating. (E) Representative DNA staining of BM MKs, BM leukocytes, and DNA staining negative control. (F) BM MK % of live, n=13 animals, 3 representative experiments. (G) Bead based BM MK quantification, n=13 animals, 3 representative experiments. (H) Representative images from imaging flow cytometry analysis of nucleated live, CD45+ lineage-CD117-CD41a+ CD42c+ cells.

**Figure 3:**
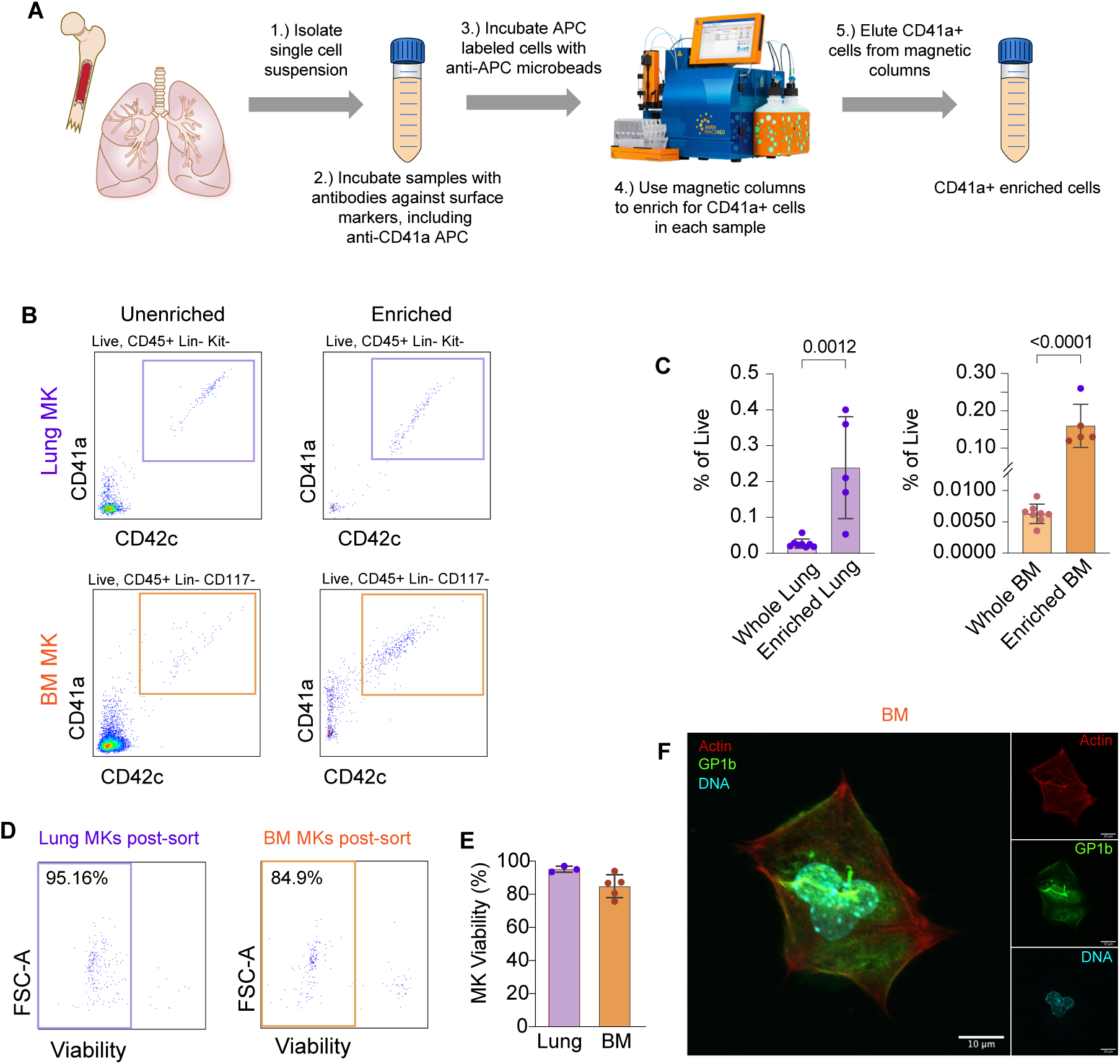
CD41 based magnetic enrichment and isolation method for lung and BM MKs (A) Experimental schematic of CD41 enrichment from lung or BM by AutoMACS. (B) Representative FACS plots of BM and lung MKs before and after enrichment. (C) BM and lung MK % of live before and after CD41 enrichment, n=13 animals, 8 individual experiments experiments, significance determined by student’s t-test. (D) Representative FACS plots of lung and BM MK post-sort viability. (E) Lung and BM MK post-sort viability analysis, n=50 animals, 5 individual experiments. (F) Representative image BM MK spread onto fibrinogen, n=20 animals, 2 individual experiments. AutoMACS Pro Separater image from Miltenyi Biotec, https://www.miltenyibiotec.com/US-en/products/automacs-pro-separator-starter-kit.html.

**Figure 4:**
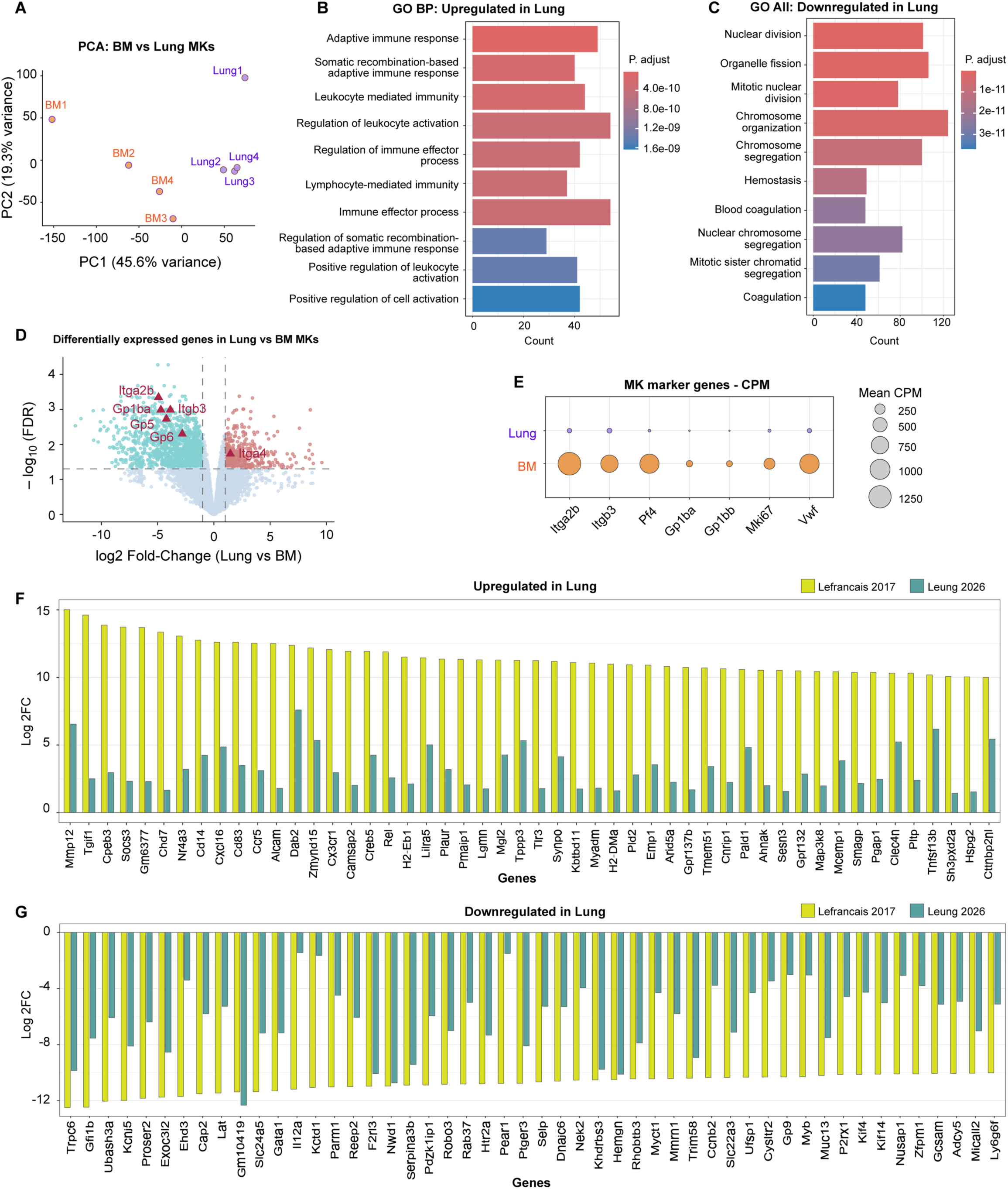
Lung MKs transcribe core MK genes while also expressing transcriptional patterns associated with MK immaturity. (A) Principal component analysis of 30,499 lung MKs and 14,232 BM MKs, 4 biological replicates each. (B) Pathways upregulated in lung MKs compared to BM MKs. (C) Pathways downpregulated in lung MKs compared to BM MKs. (D) Volcano plot depicting integrin and glycoprotein genes differentially expressed between lung and BM MKs. (E) Circle plot of notable MK genes in lung and BM MKs based on counts per million. (F) Differential expression direction and magnitude of the top 50 most downregulated genes (Log2FC) in Lefrancais, 2017, as compared to this manuscript. (G) Differential expression direction and magnitude of the top 50 most upregulated genes (Log2FC) in Lefrancais, 2017, as compared to this manuscript.

We next compared our differential expression analysis to previously published datasets comparing differential gene expression in lung MKs and BM MKs^5^. We selected the top 50 differentially expressed genes in the Lefrancais 2017 dataset and compared their direction and magnitude with the direction and magnitude of differential expression in our transcriptomic analysis (Fig4F,G). The top 50 increased genes in the Lefrancais 2017 dataset were also increased in our analysis, although the magnitude of gene expression differed, likely due to differences in cell isolation and sequencing platform. Similarly, lung MKs in both datasets had decreased expression of the same genes compared to BM MKs.

Our differential expression analysis confirmed the identity of our lung and BM MKs as characterized in our spectral and imaging flow cytometry data and agreed with differential expression patterns from the 2017 Lefrancais analysis. Furthermore our analysis supports existing data demonstrating divergent function while also suggesting lung MKs may have reduced maturity compared to BM MKs.

### Cell cycle and transcription factor (TF) genes related to MK maturity are differentially expressed in lung vs BM MKs

Considering the decreased expression of transcripts within cell cycle pathways in lung MKs, we next examined specific cell cycle genes differentially expressed in lung compared to BM MKs. The cell cycle transcripts CDC14B and PLK2, typically associated with cell cycle exit and cell stress are both more abundant in lung vs BM MKs (Fig5A)^36–40^. Additionally, CDC6, one of three factors comprising the DNA synthesis pre-replication complex (pre-RC complex), and necessary for endomitosis in MKs, is decreased in lung vs BM MKs^41–44^. Interestingly, expression of CDT1, the second of the three pre-RC complex factors, is increased in lung MKs (Fig5A).

In addition to regulating DNA replication and endomitosis, Cdc6 also serves as a transcriptional repressor of the TF c-myc. We compared differential expression of TFs, including MYC, in lung and BM MKs. Examination of prominent MK TFs showed that GATA1, GATA2, RUNX1, MYC, and ETS2 are differentially expressed between lung and BM MKs (Fig5B). Comparison of a broader set of MK-expressed TFs further highlighted the distinct transcriptional signature of lung MKs (Fig5E). Transcriptional analysis for the TFs GATA1, GATA2, and NFE2 reveal decreased expression in lung MKs compared to BM MKs (Fig5F). Gene expression analysis of the TFs RUNX1 and SRF, regulators of lineage fate and cytoskeletal development, also demonstrated significantly, but less pronounced, decreases in expression, while FLI1, another canonical MK TF, was equivalently expressed in lung and BM MKs (Fig5G). Conversely, MYC was significantly increased in lung MKs compared to BM MKs (Fig5H). Similarly, ETS2 expression was increased in lung MKs compared to BM MKs, although ETS1, while robustly expressed, was not significantly increased by FDR (Fig5H).

Immunomodulation has been implicated as the primary lung MK function, so we directly examined expression of immune genes by lung MKs in comparison to BM MKs^5, 6, 8^. Consistent with this, our data show that lung MKs have increased expression of selected anti-inflammatory genes as well as pro-inflammatory genes (Fig5C,D). IFNGR1 and IL6ST expression is increased in lung MKs, suggesting a responsiveness to interferon and IL-6 based inflammatory signaling (Fig5D). Of the anti-inflammatory genes that were increased we found the quintessential anti-inflammatory receptor IL10RA increased, as well as tumor necrosis factor (TNF) family genes TNFRSF1B, an anti-inflammatory TNF receptor, and TRAF1, downstream of TNFRSF1B (Fig5C,I). Collectively, our analysis of the lung MK transcriptome suggests a sensitivity to immunomodulatory cytokines, such as TNF, and we report for the first time a lung MK TF signature that differs from that of BM MKs.

### Lung MKs express surface integrin α4 and actively bind to fibronectin

Our transcriptional analysis identified increased expression of ITGA4 in lung MKs (Fig4D). This was of particular interest as integrin α4 is also associated with tissue homing and is expressed in the lung^45–47^. Integrin β1 can dimerize with α4, forming Very Late Antigen 4 (VLA-4), or with integrin β7 to form Lymphocyte Peyer’s patch Adhesion Molecule 1 (LPAM-1), with α4 conferring specificity to fibronectin^48, 49^. To determine whether α4 is present on lung MKs, we quantified surface expression of α4 using flow cytometry. CD117- lung and BM MKs were profiled by flow cytometry as before (Fig6A,C). Compared to controls, lung MKs express surface α4, β1, and β7 (Fig6B,D). We next hypothesized that lung MKs could bind fibronectin based on the expression of surface α4, so we incubated lung and BM MKs with labeled fibronectin and examined fibronectin binding by flow cytometry (Fig6E,F). After gating on MKs, we found fibronectin MFI on lung MKs was significantly higher than BM MKs (Fig 6G). These data show that lung MKs express cell surface α4, β1, β7, and readily bind fibronectin.

**Figure 5:**
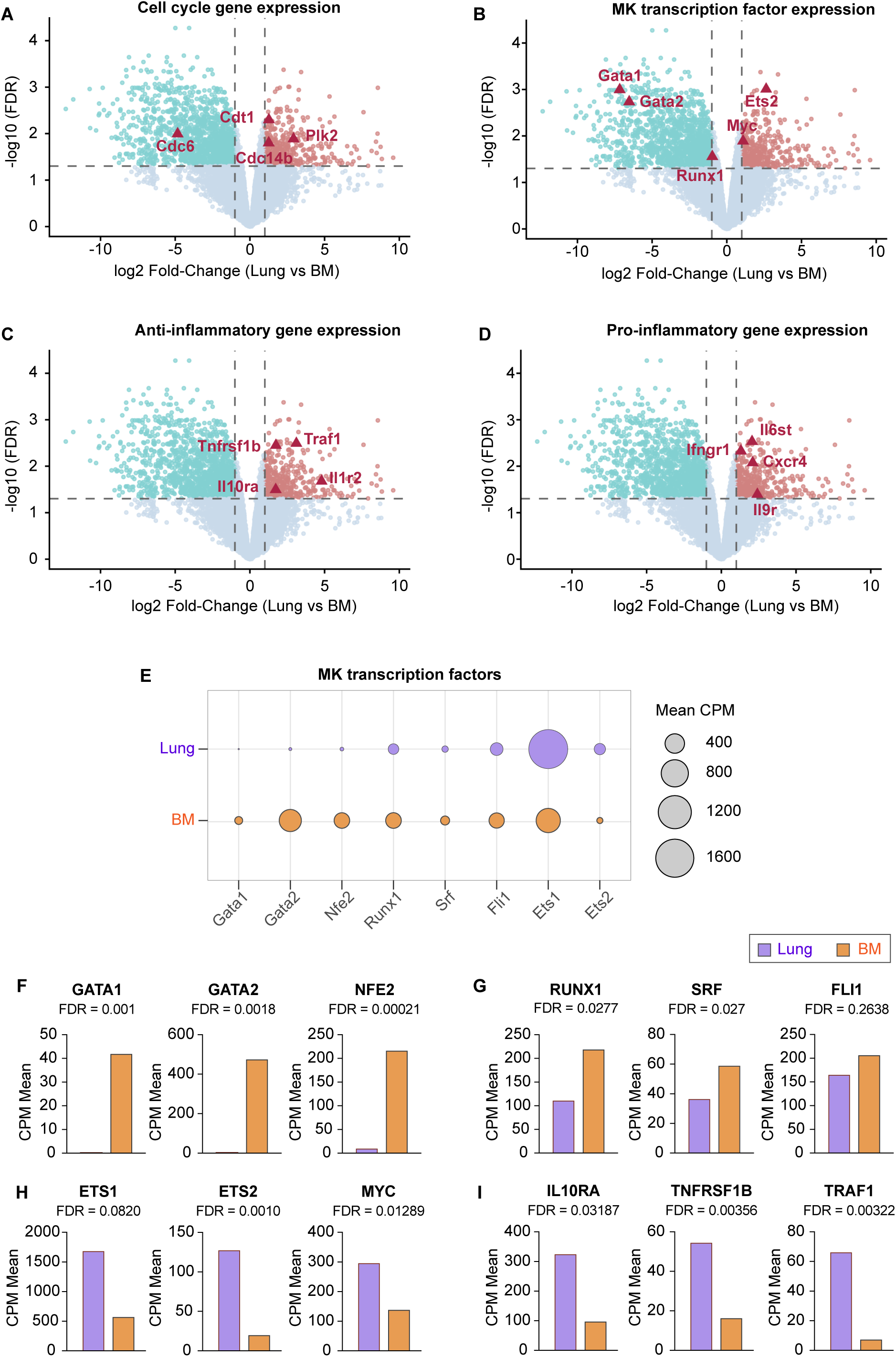
Cell cycle and transcription factor genes are differentially expressed in lung vs BM MKs. (A-D) Volcano plots depicting cell cycle, transcription factor, anti-inflammatory, and pro-inflammatory associated genes differentially expressed between lung and BM MKs. (E) Circle plot of transcription factor genes in lung and BM MKs based on counts per million (CPM). (F-H) Bar graphs of transcription factor gene expression in lung and BM MKs based on CPM, significance determined by FDR. (I) Bar graphs of immune gene expression in lung and BM MKs based on CPM, significance determined by FDR.

**Figure 6:**
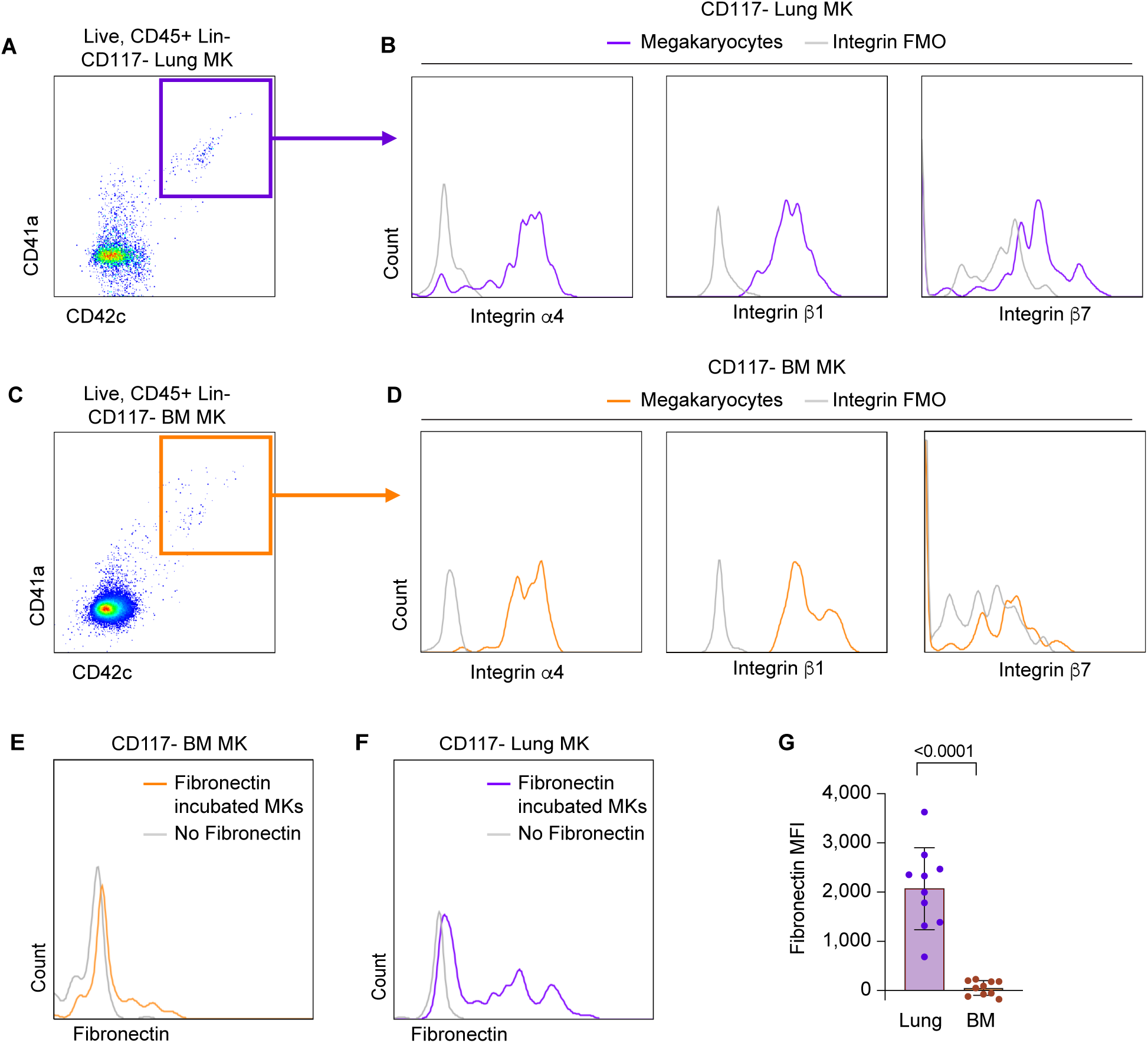
Lung MKs express surface integrin *a*4 and actively bind to fibronectin. (A) Rep FACS plot of CD117-lung MKs. (B) Representative FACS histograms of CD117-lung MK integrin *a*4, integrin β1, and integrin β7 cell surface protein expression. C) Rep FACS plot of CD117-BM MKs. (D) Representative FACS histograms of CD117-BM MK integrin *a*4, integrin β1, and integrin β7 cell surface protein expression. (E) Representative FACS histogram of labeled fibronectin bound to CD117-BM MKs. (F) Representative FACS histogram of labeled fibronectin bound to CD117-lung MKs. (G) Mean fluorescent intensity quantification of labeled fibronectin bound to CD117-BM and lung MKs. Unpaired t-test, n=20 animals, three independent experiments.

**Figure 7:**
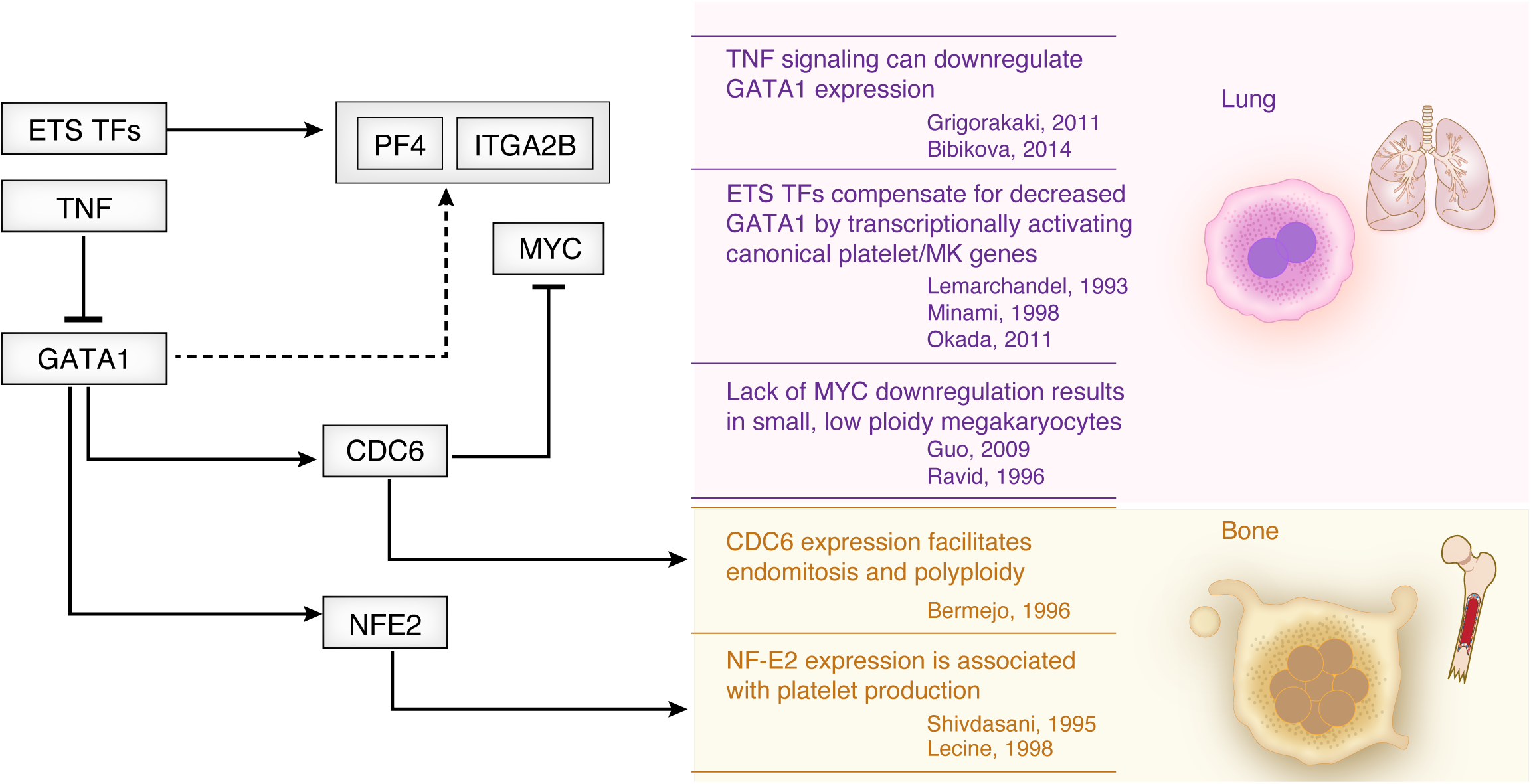
Lung and BM MK transcription factor expression may explain their divergent morphology and function. Survey of MK primary literature summarizing findings connecting MK function with TNF signaling, GATA1, ETS family transcription factors, MYC, CDC6, and NFE2.

## Discussion

Our gating scheme, isolation strategy, and transcriptomic analysis has identifed a previously unreported lung MK TF profile and ability to bind fibronectin. We validated MK identity by examining accepted patterns of MK biology, including ploidy, morphology, function, and transcriptome. Through these studies we confirm that our methods minimize the presence of leukocytes, separate progenitors, and isolate polyploid MKs that can spread onto fibrinogen coated surfaces, and express core MK genes.

A primary goal in formulating our MK characterization strategy was disaggregating developmental and leukocyte-derived heterogeneity within the MK compartment. Our MK-leukocyte analysis revealed the B-cell and myeloid markers, B220 and CD11b, were coincident with MKs (SFig1). While challenging for MK analysis, these data demonstrate the MK-leukocyte interactions that occur in the niche^10, 11, 13, 28–30^. For example, BM MKs have been reported to regulate B-cell development via production of APRIL^28^. The presence of progenitors is also a source of heterogeneity, that is not considered in standard MK characterization, despite the BM containing the entire hematopoietic tree, including CD41a+ myeloid-biased HSCs, MkPs, and MKs^18, 50,51^. By our quantifications there are approximately twice as many MkPs as MKs, and we suggest any analysis of BM MKs first involves separation of MkPs from MKs. We have determined that MKs and MkPs are 0.0055% and 0.011 % of live BM, respectively. These counts together approach those reported by Nakeff in 1974^32^. Further progress in understanding the contribution of MK subsets to health and disease depends on their disaggregation in analysis.

Our methods identify MKs with established morphology and function. Multi-nucleation is a defining MK trait, and while lung MKs may be polyploid, most reports describe 2N lung MKs^7, 8^. Our ploidy analysis shows that lung MKs are polyploid, but with decreased ploidy, suggesting lung MKs are less mature^24^. Notably, established morphological features of MKs such as large size and high ploidy are associated with maturity and platelet production, while deviations from this archetypical morphology has been associated with alternative function, such as niche regulation and immune function^5–7, 9^. Rather then a single developmental and functional progression, MK morphology may be indicative of distinct MK subsets with distinct functions, like large and small cytoplasms MKs in the BM^9^. In such a paradigm, lung MK morphology may not indicate immaturity, but rather immunomodulatory function, as has been demonstrated in the context of oropharyngeal malaria infection^5, 6, 8^.

Like lung MKs, resident tissue macrophages (RTMs) have unique niche functions, as such, they are appropriate for comparison. Alveolar macrophages, cardiac macrophages, and Langerhans cells are RTMs of the lung, heart, and epidermis, respectively, with unique tissue-specific functions^52–54^. They are identifiable by macrophage markers such as F4/80 or CD64 and perform tissue-specific functions such as clearance of excess pulmonary surfactant (alveolar macrophages), reduction of cardiomyocyte resting membrane potential (cardiac macrophages), and maintenance of skin immune homeostasis (Langerhans cells). Niche training result when primary macrophage TFs and niche activated TFs integrate to generate unique tissue-specific transcriptional programs^55–57^. Like these macrophages, the lung-resident MK transcriptome is likely influenced by signals from the lung microenvironment, such as TNF signaling.

The lung microenvironment may promote the alternative transcriptional programs found in lung MKs. Ensuing changes to MK TF expression, may then drive changes to MK morphology and function. Lung MKs express higher levels of TNFRSF1B than BM MKs, one of two TNF*a* receptors, suggesting a greater sensitivity to TNF (Fig5I). Importantly, TNF signaling has been shown to decrease GATA1 expression, which we have shown is decreased in lung MKs (Figure 5F)^58, 59^. Reduced GATA1 expression also reduces NFE2, a key platelet production TF, explaining observations of low platelet output from lung MKs at baseline (Fig5F)^8, 16, 60, 61^. ETS family TFs may compensate for reduced GATA1, as both ETS1 and GATA1 can activating MK gene transcription, such as PF4^62, 63^. Lung MKs express increased ETS2 and similar levels of ETS1 compared to BM MKs. Multiple ETS family TFs can bind to ETS sequences for PF4 activation, and both ETS2 and ETS1 can equivalently bind ETS sequences in the ITGA2B promoter^63, 64^. This alternative transcriptional activation pathway may explain decreased expression of ITGA2B, GP1BA, and PF4 in lung MKs (Fig7).

Changes to GATA1 gene expression may also affect MK morphology. Gata1 transcriptionally activates CDC6, a cell cycle mediator that functions to repress c-myc, facilitating entry into S-phase with Cdt1, and supporting endomitosis^41–43^. Reduced GATA1 and CDC6 expression may result in MKs with reduced size and ploidy (Fig5A). We did find that CDT1 is increased in lung MKs, which may partially compensate for lower levels of CDC6, resulting in some lung MK polyploidization (Figure 5A). Similarly, reduced Gata1 may also affect MK size via reduced inhibition of MYC expression^65^. MYC overexpression in MKs has been reported to result in small, proliferative, low ploidy cells^66, 67^. With reduced Gata1 in lung MKs, MYC expression remains high, preventing the development of the archetypical large, polyploid MK. The lung MK microenvironment may uniquely affect lung MK TF and cell cycle expression patterns changing resulting lung MK morphology (Fig7).

In addition to morphology, lung-specific transcriptional programs may also affect MK function. ITGA4 expression is typically associated with MK immaturity, but lung MKs express both increased ITGA4 and ITGB1, forming the homing integrin α4β1, potentially providing a mechanism for MK lung homing^34, 45, 47, 68, 69^. Surprisingly, our flow cytometric analysis showed lung MKs express more surface β7 protein then BM MKs. This analysis demonstrated that lung MKs express surface α4, β1, and β7 protein (Fig6B,D), in addition to mRNA, and suggest lung MKs may use VLA-4 or LPAM-1 signaling. Our fibronectin binding assay supported this hypothesis, showing that lung MKs also bind significantly more fibronectin then BM MKs.

Compared to archetypical BM MKs, lung MKs are smaller, lower ploidy, and functionally immunomodulatory. Here we present for the first time a characterization strategy to further dissect the biology of lung MKs, accounting for both leukocyte aggregation and developmental hierarchy. Using this strategy we have found an as yet unreported lung MK TF signature, and demonstrated for the first time that lung MKs have a greater ability to bind fibronectin then BM MKs.

## ACKNOWLEDGEMENTS

The authors thank Diana Lim and Nikita Abraham for essential graphics support, the University of Utah Comparative Medical Center for excellent animal care, and the flow cytometry and DNA sequencing core facilities for technical assistance.

## AUTHOR CONTRIBUTIONS

G.L. designed the research, performed experiments, analyzed results, interpreted data, and wrote the manuscript, M.J. analyzed results and interpreted data, M.J. performed experiments, H.B. performed experiments, I.O. performed experiments, J.R. provided critical review and edited the manuscript, M.R. provided critical review, A.C.P. designed the research and edited the manuscript, E.A.M. designed the research and edited the manuscript.

## DATA SHARING STATEMENT

For original data, please contact.

---

Supplementary Figure 1: Leukocyte marker co-stain CD41+ cells. (A) Representative FACS plots of lung CD41+ cells co-staining for F4/80, CD11b, CD3, and B220. (B) Representative FACS plots of lung CD41+ cells co-staining for CD117. (C) Percentage of lung CD41+ cells co-stained with leukocyte markers, n=10 animals, 4 independent experiments. (D) Representative FACS plots of BM CD41+ cells co-staining for F4/80, CD11b, CD3, and B220. (B) Representative FACS plots of BM CD41+ cells co-staining for CD117. (C) Percentage of BM CD41+ cells co-stained with leukocyte markers, n=11 animals, 4 independent experiments.

Supplementary Figure 2: Lung and BM MK backgating. (A) Representative FACS plots of lung CD117- and CD117+ MKs backgated onto parent gates. (B) Representative FACS plots of BM CD117- and CD117+ MKs backgated onto parent gates.

Supplementary Figure 3: Image Stream Gating Strategy: BM representative sample. (A) Representative FACS plots of image stream pre-gates, including size, singlets, DNA content, and viability (B) Representative FACS plots of image stream clean up gates, lineage exclusion, and gating on CD41a and CD42 expression. (C) Representative FACS plots of image stream gates on CD117+ or CD117- MKs. (D) Images of CD117+ MKs collected from CD117+ MK gate.

Supplementary Figure 4: CD117+ Megakaryocyte Progenitor Response to Fibrinogen. Confocal image of BM MkP spread onto fibrinogen.

Supplementary Figure 5: Quantification of MK Integrin Expression: (A) Quantification of CD117-lung MKs integrin *a*4 MFI. (B) Quantification of CD117- lung MKs integrin β1 MFI C) Quantification of CD117-MKs integrin β7 MFI. Unpaired t-test, n=24 animals, three independent experiments.

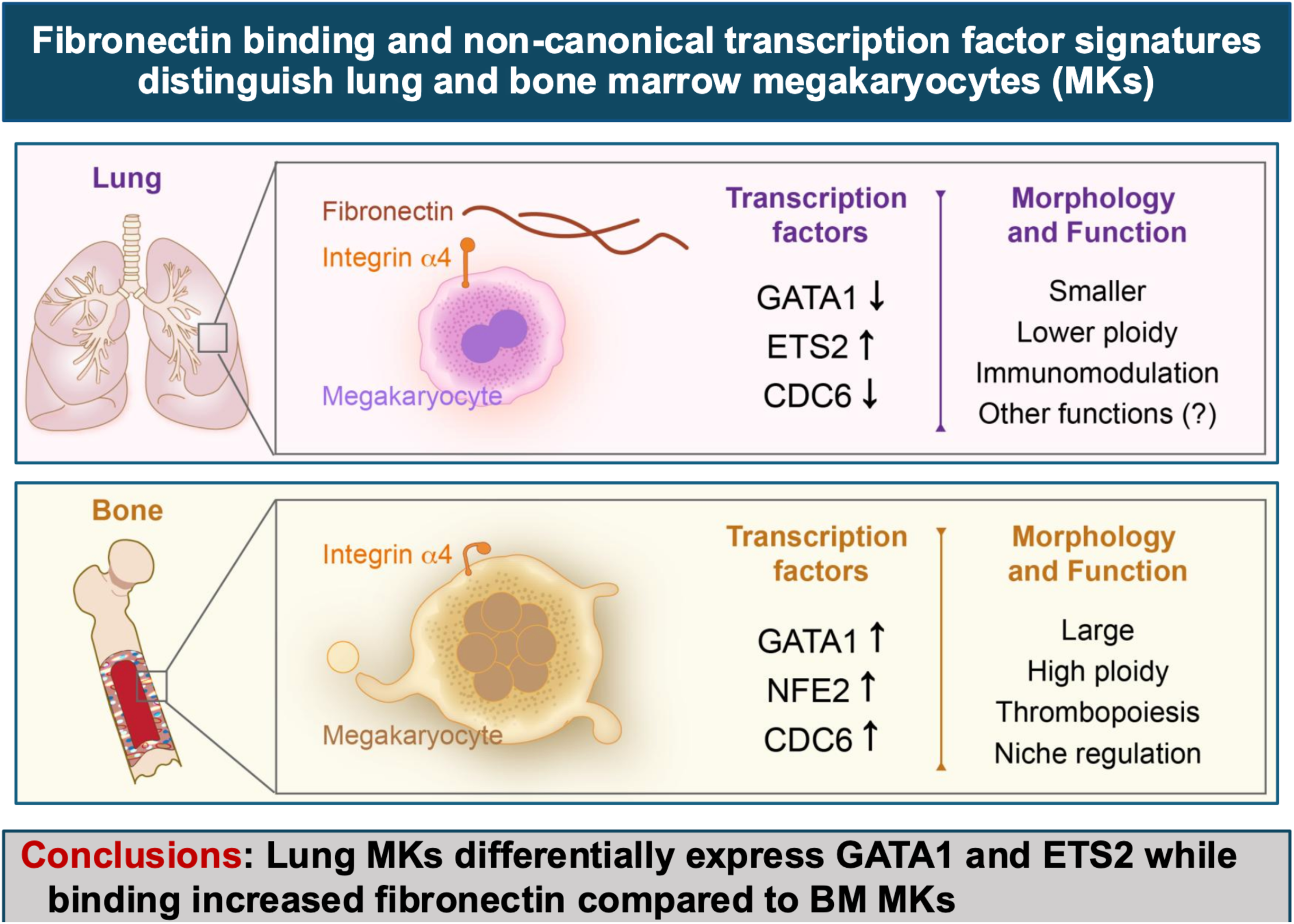

