## Supplementary Methods for "Fibronectin binding and non-canonical transcription factor signature distinguishes lung and bone marrow megakaryocytes"

Preparation of Lung homogenate

To isolate lung cells, lung tissue was removed and deposited into 3.6 mg/mL type 4 collagenase (CLS4, Worthington). Lung tissue was then chopped into pieces, and digested at 37°C for 30 minutes, while shaking at 200 RPM. After digestion, tissue was homogenized using a 16-gauge needle mounted onto a 5 mL syringe, followed by homogenization using an 18-gauge needle on a 5 mL syringe. Lung sample was then filtered through a 100-micron cell strainer and tissue was pushed through the strainer with the soft side of a 5 mL syringe plunger. Finally, the cell strainer was rinsed with 5 mM EDTA + 1x PBS. Lung sample was then centrifuged at 400 xG for 5 minutes, at 4°C. Erythrocyte lysis was performed prior to cell staining for flow cytometry analysis using ACK lysis buffer.

Erythrocyte Lysis

For flow cytometry analysis, erythrocyte lysis was performed prior to cell staining. For cell isolation, erythrocyte lysis was not performed prior to magnetic enrichment and cell staining. Erythrocyte lysis was performed by resuspending cell pellets in 0.5-1.0 mL of ACK lysis buffer (15 mM ammonium chloride, 1 mM potassium bicarbonate, and 0.01 mM disodium EDTA in distilled water, pH 7.2-7.4).

CD41-based Magnetic Enrichment

Cells were incubated with anthophycocyanin (APC) conjugated anti-CD41a for 20 minutes on ice, then washed and pelleted at 400 xG for 5 minutes. $1x10^7$ cells antibody bound cells were incubated with 20 uL anti-APC microbeads (Miltenyi catalog #130-090-855) for 20 minutes on ice, with periodic agitation by trituration every 5 minutes, then washed and pelleted at 400x G for 5 minutes. AutoMACS enrichment: positive selection mode posseld2 was used according to AutoMACS menu specifications. LS column enrichment: columns were rinsed with 1 mL of 5 mM EDTA + 1x PBS, then 4 mL of CD41 APC/anti-APC microbead conjugated cells were added to the column, which was subsequently rinsed with additional 4 mL of 5 mM EDTA + 1x PBS. Finally, cells were eluted into a tube with 4 mL of 5 mM EDTA + 1x PBS.

Flow Cytometry Bead Based Megakaryocyte Quantification

50,000 CountBrite Absolute Counting Beads (Thermofisher #C36950) were added to BM from 1 femur+tibia after centrifugation or to lung tissue from 1 lung prior to collagenase digestion, and samples were processed to single cell suspension. During flow cytometry analysis bead and cell counts were recorded. Then CD117+ or CD117- MK counts in total sample were extrapolated based on recorded bead count by multiplication of cell counts and the ratio of total/recorded beads.

$$Sample\ Count = \frac{Beads\ added}{Beads\ recorded}\ x\ Cells\ Recorded$$

Image Stream Analysis

Cells were stained with Zombie Violet (Biolegend, catalog #423113) to determine viability, then surface makers. After surface staining, cells were fixed with 2% PFA for 1 hour at room temperature. After fixation, DNA was stained with 0.1 mg/mL propidium iodide in 5 mM EDTA + 1x PBS for 20 minutes on ice. DNA stain was washed off and resuspended in 5 mM EDTA + 1x PBS for analysis. Gating strategy: Focused live singlets were identified as MKs if they were nucleated CD117- CD41+ CD42c+ lineage- cells. Resolution of lineage- cells was accomplished with the bivariate resolution afforded by plotting CD41 vs lineage and further confirmed using a lineage FMO control. Exclusion of images containing multiple cells was accomplished through a combination of aspect ratio and area, along with a nuclear spot count, ensuring that only a single nucleus was present per image. Nucleation was defined as coincidence of CD41+ morphology, CD42c+ morphology, and propidium iodide (PI) pixel intensity above 402. CD117+ status was determined using a CD117 pixel intensity threshold (63) and CD117+ colocalization with CD42c/CD41 staining; CD117+ gating was further verified using a CD117 FMO.

Fibrinogen Spreading

10,000 CD117- or CD117+ MKs were sorted from BM into MK culture media (DMEM F12, 10% FBS, 100 U/mL Pen-Strep, 50 ng/mL TPO), then cultured for 48 hours at 37°C. Glass slides were coated under sterile conditions with 250 uL of 100 ng/mL for 1 hour at 37°C, then washed three times with 1x PBS, blocked with 1% bovine serum albumin, and washed three times with 1x PBS. MKs were then sedimented onto fibrinogen-coated slides for 1 hour, then allowed to spread for 1 hour. After spreading, MKs were fixed with 4% paraformaldehyde for 1 hour, and permeabilized with 0.1% Triton X-100 for 10 minutes. MKs were visualized using immunocytochemistry staining (Supplementary Table 4) and confocal imaging (Olympus FV3000 Confocal Microscope).

RNA isolation

Cells were FACS isolated and immediately lysed using Trizol (Invitrogen #15596026). For RNA isolation, RNA samples were pre-incubated with 1 µl 20 mg/ml glycogen, then the Trizol protocol was used. After RNA isolation, samples were DNase treated (ThermoFisher #AM1907) and cleaned (Zymo RCC-5).

Bioinformatic processing

Raw paired-end RNA-seq reads were processed with the nf-core/rnaseq pipeline v3.21.0[25] Reads were aligned to the mouse reference genome GRCm39 (GENCODE M38, Ensembl release 115) using STAR, followed by transcript-level quantification with Salmon. UMI deduplication was performed with UMI-tools.

Differential expression analysis

Principal component analysis (PCA) was performed on log2-CPM-transformed, mean-centred and scaled expression values across all retained genes. Sample-level clustering was visualised with a row-Z-scored heatmap of the top 1,000 most variable genes (ranked by row variance of log2 CPM) using pheatmap with hierarchical clustering on both axes. Gene-level counts for bone marrow megakaryocytes (BM MK, n = 4) and lung megakaryocytes (Lung MK, n = 4) were analysed in R v4.3.1 using edgeR v3.42.4[26]. Library sizes were normalised by the trimmed mean of M-values (TMM) method[27]. Genes were retained if their mean CPM exceeded 2 in at least one of the two groups (13,702 of 78,334 genes retained). Gene-wise dispersions were estimated with estimateDisp(robust = TRUE), and differential expression was tested using the quasi-likelihood F-test (glmQLFit/glmQLFTest), contrasting Lung vs BM. P-values were adjusted by the Benjamini–Hochberg procedure; genes with FDR < 0.05 and |log2FC| > 1 were considered differentially expressed. FPKM values were computed from TMM-normalised effective library sizes and union exonic gene lengths derived from the project GTF.

Gene Ontology enrichment

Over-representation analysis of Gene Ontology terms was performed with clusterProfiler v4.8.3[28] using enrichGO() against the org.Mm.eg.db annotation database. Enrichment was tested separately for genes upregulated in Lung MKs and BM MKs (FDR < 0.05, |log2FC| > 1), for the Biological Process ontology and for all ontologies pooled. P-values were adjusted by the Benjamini–Hochberg procedure with a q-value cutoff of 0.2.

Software and packages

Analyses were performed in R v4.3.1. Bioconductor and CRAN packages used: edgeR v3.42.4, GenomicFeatures, clusterProfiler, org.Mm.eg.db, AnnotationDbi, dplyr, tidyr, ggplot2 v4.0.2, ggrepel, pheatmap, RColorBrewer, and openxlsx. Upstream processing used nf-core/rnaseq v3.21.0 (STAR, Salmon, UMI-tools).
