## Supplementary Tables for "Fibronectin binding and non-canonical transcription factor signature distinguishes lung and bone marrow megakaryocytes"

| Table 1 - Leukocyte and Lymphocyte Antibodies | | |
|---|---|---|
| Marker | Antigen/Clone | Cell Type Specificity |
| F4/80 | BM8 | Macrophages |
| FcER1a | MAR1 | Mast Cells |
| CD11b | M1/70 | Myeloid Cells |
| Ly6c | HK1.4 | Monocytes |
| Ly6g | 1A8 | Neutrophils |
| Ter119 | TER119 | Erythrocytes |
| B220 | RA3-6B2 | B-cells |
| CD3 | 145-2C11 | T-cells |

| Table 2 - Leukocyte Antibodies | | |
|---|---|---|
| Marker | Clone | Conjugation |
| F4/80 | BM8 | Biotin |
| FcER1a | MAR1 | Biotin |
| CD3 | 145-2C11 | Biotin |
| B220 | RA3-6B2 | Biotin |
| Ter119 | TER119 | Biotin |
| NK1.1 | PK136 | Biotin |
| CD11b | M1/70 | Biotin |
| Gr-1 | RB6-8C5 | Biotin |

| Table 3 - MK Antibodies | | |
|---|---|---|
| Marker | Clone | Conjugation |
| CD45 | 30-F11 | BUV395 |
| CD117 | ACK2 | BV605 |
| CD150 | TC15-12F12.2 | BV785 |
| Sca-1 | E13-161.7 | PB |
| CD41a | MWReg30 | PE-Cy7 |
| CD42c | X488 | Alexa Fluor 488 |

| Table 4 - Immunocytochemistry Stain | | |
|---|---|---|
| Reagent | Staining Target | Excitation/Emission (nm) |
| DAPI | DNA | 355/405 |
| X488 | GP1b beta | 488/488 |
| Actin Red | F-Actin | 540/565 |

| Table 5 - MK Integrin Antibodies | | |
|---|---|---|
| Reagent | Clone | Conjugation |
| CD49d | R1-2 | Real Blue 780 |
| CD29 | HM β1-1 | Real Blue 670 |
| ITGB7 | M293 | Real Yellow 655 |
